# Novel EGFR-Mutant Mouse Models of Lung Adenocarcinoma Reveal Adaptive Immunity Requirement for Durable Osimertinib Response

**DOI:** 10.1101/2022.10.26.513856

**Authors:** Emily K Kleczko, Anh T Le, Trista K Hinz, Teresa T Nguyen, Andre Navarro, Cheng-Jun Hu, Eric T Clambey, Daniel T Merrick, Mary Weiser-Evans, Raphael A Nemenoff, Lynn E Heasley

**Affiliations:** Departments of Medicine, University of Colorado Anschutz Medical Campus, Aurora, CO; Departments of Craniofacial Biology, University of Colorado Anschutz Medical Campus, Aurora, CO; Departments of Anesthesiology, University of Colorado Anschutz Medical Campus, Aurora, CO; Departments of Pathology, University of Colorado Anschutz Medical Campus, Aurora, CO; Eastern Colorado VA Healthcare System, Rocky Mountain Regional VA Medical Center, Aurora, CO

**Keywords:** Lung adenocarcinoma, EGFR, GEMM, adaptive immunity, tyrosine kinase inhibitor

## Abstract

Lung cancers bearing oncogenically-mutated EGFR represent a significant fraction of lung adenocarcinomas (LUADs) for which EGFR-targeting tyrosine kinase inhibitors (TKIs) provide a highly effective therapeutic approach. However, these lung cancers eventually acquire resistance and undergo progression within a characteristically broad treatment duration range. Our previous study of EGFR mutant lung cancer biopsies highlighted the positive association of a TKI-induced interferon γ transcriptional response with increased time to treatment progression. To test the hypothesis that host immunity contributes to the TKI response, we developed novel genetically-engineered mouse models of EGFR mutant lung cancer bearing exon 19 deletions (del19) or the L860R missense mutation. Both oncogenic EGFR mouse models developed multifocal LUADs from which transplantable cancer cell lines sensitive to the EGFR-specific TKIs, gefitinib and osimertinib, were derived. When propagated orthotopically in the left lungs of syngeneic C57BL/6 mice, deep and durable shrinkage of the cell line-derived tumors was observed in response to daily treatment with osimertinib. By contrast, orthotopic tumors propagated in immune deficient *nu/nu* mice exhibited modest tumor shrinkage followed by rapid progression on continuous osimertinib treatment. Importantly, osimertinib treatment significantly increased intratumoral CD3+ T cell content relative to diluent treatment. The findings provide strong evidence supporting the requirement for adaptive immunity in the durable therapeutic control of EGFR mutant lung cancer.

## Introduction

The epidermal growth factor receptor (EGFR) is mutated in 15-30% of lung adenocarcinomas (LUADs) [[1-4] where the L858R missense mutations and in-frame exon 19 deletions account for ∼85% of EGFR-activating mutations [3]. The EGFR tyrosine kinase inhibitor (TKI) erlotinib was approved in 2004 for non-small cell lung cancer (NSCLC) patients [5]. Subsequently, it was noted that novel EGFR mutations predicted patient sensitivity to erlotinib [6, 7], and EGFR-specific TKIs became first-line therapies for LUAD patients whose tumors harbor EGFR mutations [5, 8, 9]. To combat the EGFR-T790M mutation that arose as a resistance mechanism in the setting of 1^st^ generation TKIs, a 3^rd^ generation inhibitor, osimertinib, that inhibits oncogenic EGFR with or without the T790M mutation [10-12] was approved as the present first-line therapy for EGFR mutant LUAD. Despite these important advances in precision oncology for oncogenic EGFR-driven LUAD, the vast majority of patients will develop acquired resistance and eventually relapse [13, 14]. There is presently no FDA-approved therapy for patients progressing on osimertinib.

While oncogenic EGFR mutations predict responsiveness to EGFR-specific TKIs[6, 7, 15], patients display wide-ranging therapeutic responses with rare complete responses, but largely partial responses accompanied by residual disease [16, 17]. Thus, defining molecular mechanisms contributing to residual disease and variable duration of EGFR TKI therapeutic responses may provide opportunities for novel drug combinations to improve outcomes for this oncogene-defined group of patients. Based on their lack of response to immunotherapies and lower mutation burden, it is tempting to consider host immunity as irrelevant to EGFR-driven LUAD subsets and therapeutic response to TKI. However, we recently reported that the degree of induction of an interferon γ (IFNγ) transcriptional response in “on-treatment” biopsies obtained from EGFR mutant LUAD patients positively associated with the duration of the therapeutic response [18]. This same study and that of Maynard et al [19] provided evidence for increased tumoral content of T cells in response to TKI treatment, supporting a growing body of evidence that host immunity contributes to the therapeutic response to oncogene-targeted agents [20-28]. While human tumor specimens have provided important associations, relevant preclinical models are necessary to provide deeper mechanistic insight into tumor-immune cell interactions.

There are limited preclinical models available to explore how innate and adaptive immunity may contribute to the TKI response in EGFR-mutant lung tumors. Human patient-derived EGFR mutant lung cancer cell lines provide models for assessing direct actions of TKIs on tumor cells, but *in vivo* studies require implantation into immune-deficient mice which lack T and B cells. Two distinct groups have developed genetically-engineered mouse models (GEMMs) where human EGFR cDNAs bearing L858R or exon 19 deletions are expressed in relevant lung cell types via tetracycline-inducible promoters to yield multi-focal lung tumors [29-33]. These models have provided clear evidence for the therapeutic activity of TKIs and antibody-based agents, but due to the selection of the tetracycline-inducible system, cultured cell lines have not been generated. In this study, we developed novel GEMMs whereby murine EGFR cDNAs encoding the two major oncogenic mutations are expressed following Cre recombinase-mediated excision of Lox-stop-Lox sequences, yielding multifocal lung tumors in mice. Our group has established an orthotopic model whereby murine lung cancer cells are implanted directly into the left lung of syngeneic mice, resulting in the development of a single tumor in the appropriate microenvironment complete with a fully-competent immune system [34-36]. This model is well suited for examining interactions between the cancer cells and the tumor microenvironment (TME), and has allowed us to define pathways in the cancer cells associated with response to immunotherapy [37, 38]. Using these oncogenic EGFR GEMMs, we have established transplantable murine lung cancer cell lines that yield orthotopic lung tumors suitable for assessing the role of host immunity in the therapeutic response to TKI. The results reveal that engagement of adaptive immunity is necessary for durable *in vivo* efficacy of osimertinib in these models of EGFR-mutant LUAD.

## Materials and Methods

### Generation of Mouse Models of EGFR-Mutant LUAD

The murine *Egfr* cDNA was amplified from reverse-transcribed mRNA isolated from the Lewis Lung Cancer (LLC) cell line with forward primer 5’-ACCGCTAGCGCCGCCACC**ATG**CGACCCTCAGGGACCGC-3’ and reverse primer 5’-ACCAAGCTT**TCA**TGCTCCAATAAACTCACT-3’. The bolded ATG and TCA represent the translation start and stop sites, respectively of murine *Egfr*. The purified 3600 bp amplicon was cloned into pcDNA3.1/Hygro (Invitrogen, Waltham, MA) and validated by Sanger sequencing. Murine *Egfr* cDNAs encoding the L860R mutation (equivalent to human L858R) and EGFR-ELREA (aa 748-752) deletion (del19) were generated by *in vitro* PCR-based mutagenesis. Egfr-transgenic mice (Rosa-LoxP-STOP-LoxP-EGFR-L860R/EGFR-del19-mC3-WPRE) were generated by the Mouse Genetics Core Facility at National Jewish Health (Denver, CO). Founding mice were subsequently bred with Trp53^LoxP^ mice (B6.129P2-*Trp53*^*tm1Brn*^/J; Jackson Laboratory stock #008462). Intratracheal administration of Adeno-Cre virus (2.5 × 10^7^ PFU/mouse) led to the development of multifocal tumors within 8 weeks (see below).

### Genotyping

DNA was isolated from mouse ear clips following the Extracta DNA Prep for PCR (Quantabio #95091) following the manufacturer’s protocol. PCR for the gene of interest was performed using the AccuStart II GelTrack PCR SuperMix protocol (Quantabio, #95136, Beverly, MA).

<u>EGFR</u> – Forward primer: 5’-AGTTGTTATCAGTAAGGGAGCTGCA-3’; Reverse primer: 5’-ACCGAAAATCTGTGGGAAGTCTTGT-3’; RS-CAG rev1 primer: 5’-CTCGACCATGGTAATAGCGATGAC-3’; PCR Program: 95°C 3 min, [95°C 30 sec, 58°C 30 sec, 72°C 40 sec] x 38 cycles, 72°C 3 min, 4°C 5 min.

<u>TP53</u> [39] – Forward primer: 5’-GGTTAAACCCAGCTT GAC CA-3’; Reverse primer: 5’-GGAGGCAGAGACAGTTGGAG-3’; PCR Program: 94°C 1 min, [94°C 30sec, 60°C 30 sec, 72°C 1 min] x 35 cycles, 72°C 10 min, 4°C 5 min.

### Generation and Maintenance of EGFR-Mutant Murine Lung Cancer Cell Lines

To generate mEGFR-del19.1 and mEGFR-L860R.1 cells, lungs from AdCre-infected EGFR-transgenic mice were harvested and tumors minced, digested and cultured. For the generation of mEGFR-del19.2 cells, a tumor from an EGFR-transgenic mouse was minced into 1 mm^3^ pieces, dipped in Matrigel (Corning, #354234, Corning, NY), and implanted into the flanks of a *nu/nu* mouse. The resulting flank tumor was harvested, minced, digested and cultured on a plastic tissue culture dish in Roswell Park Memorial Institute (RPMI)-1640 medium containing 10% fetal bovine serum. EGFR-mutant murine cell lines were passaged in culture until stable epithelial cell lines were established. The presence of the predicted EGFR mutation was verified by Sanger Sequencing of the EGFR-mutant transgene. Cells were cultured and maintained at 37°C in RPMI-1640 media (Corning) with 10% fetal bovine serum (FBS; Gibco, Waltham, MA) and 1% penicillin/streptomycin (Corning) in a humidified incubator with 5% CO_2_. Cells were passaged for no more than 10 passages before a new vial of cells was thawed.

### In vivo Animal Studies

C57BL/6J mice were purchased from Jackson Laboratory (#000664), and *nu/nu* mice (Hsd:Athymic Nude-*Foxn1*^*nu*^; #069) were obtained from Envigo. Transgenic mice were developed at the Mouse Genetics Core at National Jewish Hospital. Experiments were performed on 8 to 12 week-old male and female mice that were bred (transgenic EGFR mice), housed, and maintained at the University of Colorado Anschutz Medical Campus vivarium, and all procedures and manipulations were performed under an Institutional Animal Care and Use Committee-approved protocol. Mice were sacrificed using CO_2_ and cervical dislocation as a secondary method.

#### Orthotopic Lung Model

We have previously reported our orthotopic lung injection model [35, 36, 40]. Briefly, 5 × 10^5^ murine EGFR-mutant lung cancer cells in 40 μL of 1.35 mg/mL Matrigel diluted in Hank’s Balanced Salt Solution (HBSS; Corning) were injected into the left lobe of the lungs of either C57BL/6 or *nu/nu* mice. For the injection, mice were anesthetized using isoflurane, their left side was shaved, a 1 mm incision was made in the skin on their left side to visualize the ribs and the left lung, cells were injected using a 30-gauge needle, and the incision was closed with staples. Mice were imaged 7-10 days post injection for a baseline pre-treatment image, and then randomized into groups (n=10) treated with osimertinib (5 mg/kg; MedChemExpress, Monmouth Junction, NJ) or control (H_2_O) by oral gavage with a schedule of 5 days on treatment followed by 2 days off. For immunofluorescence experiments, tumors were allowed to establish for 3 weeks and then treated by oral gavage for 4 days with 5 mg/kg osimertinib or control. Mice were euthanized, tumors measured via digital calipers, and tissue was harvested (see below).

#### Xenograft Flank Model

1 × 10^6^ cells in 100uL of 1.35 mg/mL matrigel diluted in PBS were injected into each flank of *nu/nu* mice. When the tumors reached ∼200mm^3^, mice were randomized to diluent control (H_2_O) or 5 mg/kg osimertinib delivered by daily oral gavage. Tumor volumes were measured every 3 days via digital calipers. Mice were euthanized when tumor size reached ∼1000 mm^3^ per IACUC guidelines.

### Micro-CT (μCT) Imaging

μCT imaging was performed by the Small-Animal IGRT Core at the University of Colorado Anschutz Medical Campus in Aurora, CO using the X-Rad 225Cx Micro IGRT and SmART Systems (Precision X-Ray, Madison, CT). Tumor volume was quantified from μCT images using ITK-SNAP software [41] (www.itksnap.org).

### Tissue Harvesting and Processing for Immunofluorescence

For formalin-fixed paraffin-embedded (FFPE) tissue, the lungs of mice were perfused with 5mL of PBS/heparin (20 U/mL; Sigma) and inflated with 4mL of 4% paraformaldehyde (PFA; Electron Microscopy Scientific). The tumor-bearing left lung was fixed in 4% PFA for 24 hours, upon which it was switched to 70% ethanol. Tissues were processed and embedded into FFPE by the Pathology Shared Resource at the University of Colorado Anschutz Medical Campus. The Pathology Shared Resource cut blank slides performed hematoxylin and eosin staining (H&E). Paraffin embedded samples were cut into 5 μm sections. Sections were dehydrated, immersed in 0.1% Sudan Black B (Sigma, #199664-25G) in 70% ethanol for 20 minutes, washed in TBST, incubated in citrate antigen retrieval solution at 100°C for 2 hours, washed in 0.1M glycine/TBST (Sigma, #G-8898) for 10 minutes, and placed in 10 mg/ml sodium borohydride in Hank’s buffer (Gibco, #14175-095). After blocking with 10% goat serum in equal parts of 5% BSA in TBST and Superblock (ScyTek Laboratories, #AAA999, Logan, UT) at 4°C overnight, sections were incubated with primary anti-CD3 (Thermo Scientific, #MA5-14524, Waltham, MA) in equal part solution of 5% BSA in TBST and Superblock for 1 hour at room temperature, washed in TBST, and incubated with secondary goat anti-rabbit IgG Alexa Flour 488 (Invitrogen, #A11034). Slides were mounted with Vectashield mounting medium with DAPI (Vector Laboratories, #H-1200, Newark, CA). Positive staining was determined by count per high power field (40X), each tumor count was an average of 3 different fields per tumor, of 3-4 distinct tumors, and averaged between two different observers.

### RNAseq

Murine EGFR mutant cell lines cultured in 10 cm dishes were treated for 1-6 days with 0.1% DMSO or 100 nM osimertinib. RNA was submitted to the University of Colorado Cancer Center Genomics Shared Resource where libraries were generated and sequenced on the NovaSeq 6000 to generate 2×151 bp reads. Fastq files were quality checked with FastQC, Illumina adapters trimmed with bbduk, and mapped to the mouse mm10 genome with STAR aligner. Counts were generated by STAR’s internal counter and reads were normalized to counts per million (CPM) using the edgeR R package (20). Heatmaps were generated in Prism 9 (GraphPad Software, San Diego, CA).

### Immunoblotting

Cells were collected in phosphate-buffered saline, centrifuged, and suspended in lysis buffer (0.5% Triton X-100, 50 mM β-glycerophosphate (pH 7.2), 0.1 mM Na_3_VO_4_, 2 mM MgCl_2_, 1 mM EGTA, 1 mM DTT, 0.3 M NaCl, 2 μg/ml leupeptin and 4 μg/ml aprotinin). Aliquots of the cell lysates containing 50 μg of protein were submitted to SDS-PAGE and immunoblotted for proteins described in the legend to Figure 3.

### Proliferation Assays

The murine EGFR mutant cell lines were plated at 100 cells per well in 96-well tissue culture plates. Twenty-four hours later, cells were treated in triplicate with the indicated concentration of gefitinib or osimertinib. Cell number assessed by DNA content was determined after 7-10 days of culture using CyQUANT Direct Cell Proliferation Assay (Life Technologies, #C35011, Carlsbad, CA) according to manufacturer’s instructions. Data are presented as percent of control cells treated with 0.1% DMSO.

### Statistics

All graphing and statistical analyses were performed using GraphPad Prism version 9.2.0. Data are presented as the mean ± standard error of the mean (SEM).

## Results

### Transcriptional induction of gene signatures associated with immune engagement in biopsies from TKI-treated patients

We previously performed RNAseq analysis on 8 pairs of patient-derived EGFR mutant lung tumor biopsy specimens obtained prior to and after ∼2 weeks of treatment with an EGFR-specific TKI [18]. The findings revealed induction of multiple immune and inflammation-related pathways in “on-treatment” relative to pre-treatment biopsies, and the degree of induction of the IFNγ Hallmark pathway was positively associated with the time to progression (TTP) experienced by the patients. The RNAseq data were used to calculate scores from the T cell-inflamed gene expression profile reported by Ayers et al [42] as well as a cytotoxic T cell signature developed by Lau et al [43]. Both of these signatures were found to associate with response to anti-PD-1 immunotherapy. The findings reveal that the Ayers signature scores increased in all on-treatment biopsies from patients with TTP of greater than 12 months (p < 0.05 by paired t-test), but in only 1 patient with a TTP less than 12 months (**Fig.1A**). Also, the Lau et al signature (**Fig. 1B**) was increased in all four patients with TTP greater than 12 months and trended towards significance (p = 0.065). Thus, the duration of TKI response in this set of patients associated with induction of these gene signatures despite the fact that EGFR mutant lung cancers fail to exhibit significant response to anti-PD-1-based immune therapies [44-46]. We interpret the increased induction of the IFNγ Hallmark pathway as well as the aforementioned gene signatures in **Fig. 1** as evidence supporting the contribution of host T cells to the therapeutic response of EGFR mutant lung cancer to TKIs. Definitively testing this hypothesis requires experiments with EGFR-driven lung cancer models in fully immune competent hosts.

**Figure 1.**
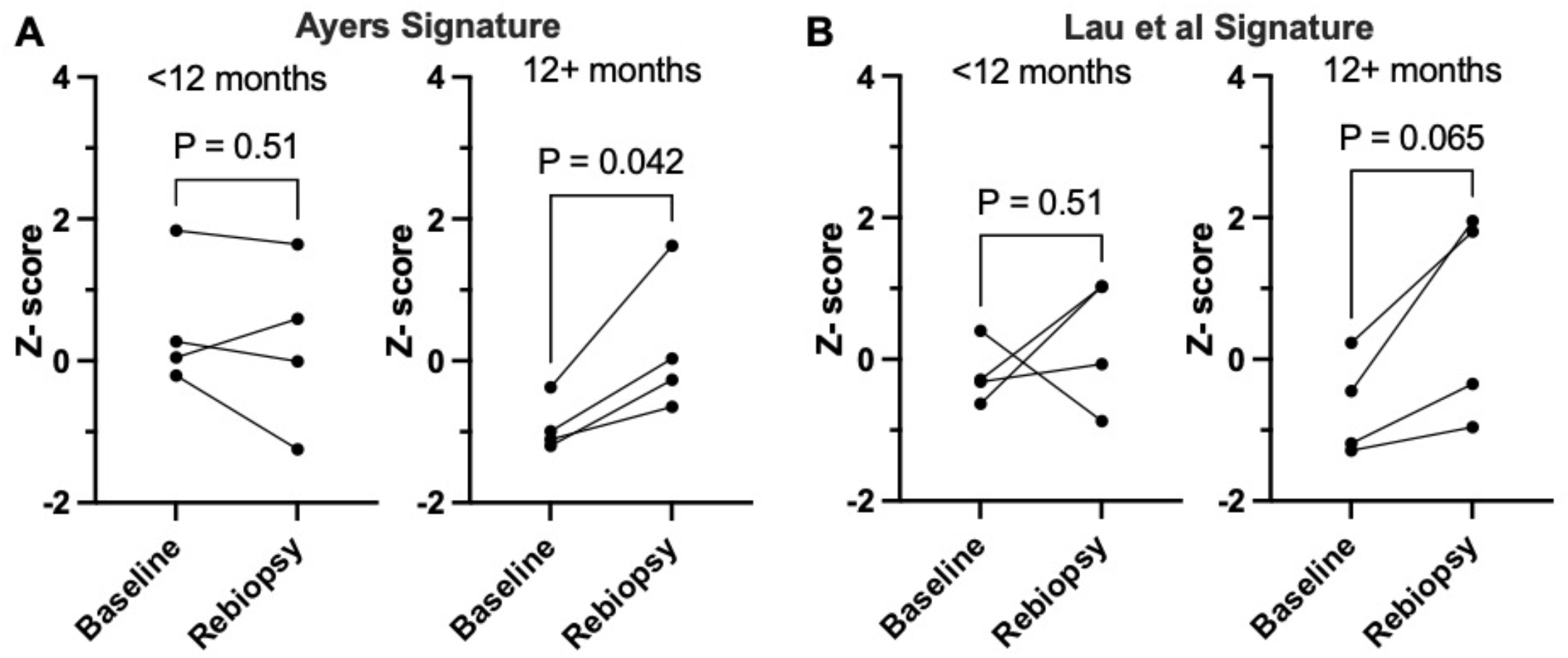
Induction of T cell-focused gene expression signatures in on-treatment biopsies from EGFR mutant LUAD patients associates with greater time to progression. Using previously reported RNAseq data from pre- and on-treatment biopsies collected from EGFR mutant LUAD patients, the sums of the 18 genes in the signature described by **(A)** Ayers et al [42] and the 25 genes in the signature described by **(B)** Lau et al [43] were calculated for each patient-derived specimen and converted to Z-scores. The resulting values for the Ayers (**A**) and Lau et al (**B**) signatures were binned by TTP of less than or greater than 12 months where the eight patients exhibited TTP of 6, 6.2, 8.3, 8.6 12.5, 13, 13.1 and 16.3 months (mean and median = 10.5 and 10.6 months, respectively). The matched Z-scores for each patient at baseline and upon re-biopsy are graphed and were analyzed by paired t-tests. The p values for the two analyses are indicated.

### Generation of mouse models that develop EGFR mutation-driven LUAD

To provide preclinical models of EGFR-mutant LUAD from which murine LUAD cell lines could be established, novel EGFR-mutant GEMMs were developed. The murine *Egfr* cDNA was amplified from reverse-transcribed RNA prepared from murine Lewis Lung carcinoma (LLC) cells and the exon 19 deletion and L860R (equivalent to human EGFR L858R) mutations were generated by PCR-directed mutagenesis (see Materials and Methods). Transgenic mouse strains were generated whereby the *Efgr*^del19^ or *Egfr*^L860R^ transgenes were inserted into the Rosa locus controlled by a strong CAG promoter and a proximal LoxP-STOP-LoxP sequence (**Fig. 2A**). After interbreeding with Trp53^LoxP^ mice, adeno-Cre virus preparations (AdCre) were administered intratracheally (see Materials and Methods) and 8-12 weeks later, multifocal lung adenocarcinomas developed with some mice exhibiting tumor formation as early as 4 weeks post AdCre injection (**Suppl. Table S1**). The tumors were readily visualized by gross inspection (**Fig. 2B**) and pathological analysis of H&E-stained sections identified hyperplasias, atypical adenomatous hyperplasias (AAH), adenomas, and adenocarcinomas (**Figure 2C** and **Suppl. Table S1**). To test the dependency of the tumors arising in these models on EGFR signaling, an Efgr^del19^ mouse intratracheally injected with AdCre was imaged 15 weeks post-injection by *μ*CT, and then treated daily with osimertinib by oral gavage. Osimertinib treatment of the mouse for 3 weeks induced significant shrinkage of tumors detected by *μ*CT (**Suppl. Fig. S1**).

**Figure 2.**
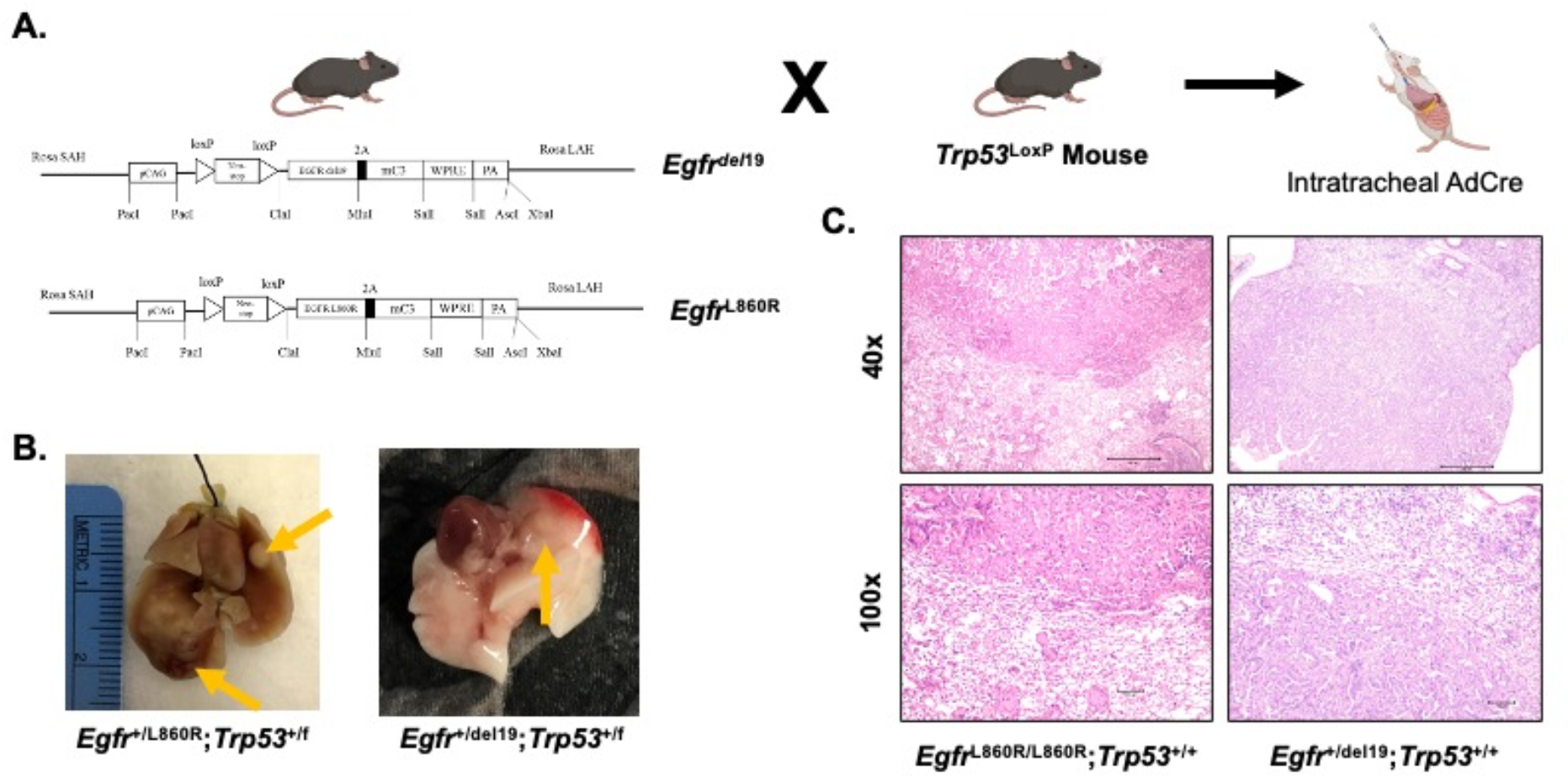
Transgenic mouse strains for *Cre*-mediated expression of *Egfr*^del19^ and *Egfr*^L860R^. *Egfr*-transgenic mice (Rosa-loxP-STOP-LoxP-EGFR L860R/del19-mC3-WPRE) were created by the Mouse Genetics Core at National Jewish Health. One strain expresses the murine Egfr cDNA encoding an exon 19 deletion and one with an L860R mutation (equivalent to the human L858R mutation). The resulting mice were bred with p53^*LoxP*^ mice from Jackson Laboratories (B6.129P2-*Trp53*^*tm1Brn*^/J; #008462). To induce expression of the oncogenic EGFR transgene, an Adeno-Cre (AdCre) virus was instilled intratracheally (2.5×10^7^ PFU/mouse) to excise the LoxP-STOP-LoxP sequence. **(A)** Schematic of the recombined Rosa locus and subsequent breeding crosses. **(B)** Gross dissection of multifocal tumors in the right and left lungs resulting from AdCre instillation. Left panel: *Egfr*^+/L860R^;*Trp53*^+/f^ lungs at 24 weeks post AdCre. Right panel: *Egfr*^+/del19^;*Trp53*^+/f^ lungs at 11 weeks post AdCre. Arrows indicate visible tumors. **(C)** H&E-stained tumor sections showing adenocarcinoma of the lungs in this model resulting from AdCre installation. Left panel: an *Egfr*^L860R/L860R^;*Trp53*^+/+^ tumor at 14 weeks post AdCre. Right panel: an *Egfr*^+/del19^;*Trp53*^+/+^ tumor at 14 weeks post AdCre. Images captured at 40x and 100x; Scale bar = 500 μm for 40x images and 100 μm for 100x images.

### Generation of murine EGFR-mutant lung cancer cell lines

To generate cell lines from lung tumors that arise in the Efgr^del19^ and Egfr^L860R^ GEMMs, individual tumors were dissected from the lungs of mice previously injected with AdCre, dissociated into single cell suspensions and cultured until stable cell lines were established. Two cell lines were generated by direct propagation on tissue culture plastic, mEGFRdel19.1 and mEGFR-L860R.1 (**Suppl. Fig. S2A**). For a third cell line (mEGFRdel19.2), a primary tumor was minced and implanted in the flank of a *nu/nu* mouse. The resulting flank tumor was harvested, dissociated, and cultured on plastic until a cell line was established (**Suppl. Fig. S2A**). The transgene construct also encodes cerulean blue, and fluorescence microscopy confirmed expression of this gene in mEGFRdel19.2 cells (**Suppl. Fig. S2B**). Expression of the oncogenic EGFR cDNAs was confirmed in all cell lines by direct Sanger sequencing (**Suppl. Fig. S2C**). The three murine EGFR cell lines were confirmed to be p53-null as assessed by immunoblot analysis, including mEGFRdel19.1 which was established from a p53^+/f^ mouse (**Suppl. Fig. S2D**).

### *In vitro* and *in vivo* sensitivity of murine EGFR mutant lung cancer cell lines to TKIs

The growth and MAPK signaling dependency of the murine cell lines on mutant EGFR was tested *in vitro*. Measurement of cell growth in the presence of increasing concentrations of gefitinib and osimertinib revealed similar dose-dependent growth inhibition in the three cell lines with IC_50_ values in the 3-10 nM range for both drugs (**Fig. 3A**). Treatment of the cell lines with EGFR-targeting TKIs (gefitinib, afatinib, osimertinib) but not the ALK/MET-targeting TKI, crizotinib inhibited phospho-Y1068 content of EGFR as well as levels of pS483-AKT, and pThr202/Tyr204-ERK1/2 (**Fig. 3B-D**). The findings demonstrate that the murine EGFR lung cancer cell lines are dependent on oncogenic EGFR for growth as well as MAPK and AKT signaling.

An orthotopic mouse model of lung cancer whereby murine lung cancer cells are directly inoculated into the left lobe of C57BL/6 mice [[35, 36, 40] was deployed to assess the therapeutic response of these murine EGFR mutant lung cancer cell lines in the context of an immune-competent TME. Both mEGFRdel19.1 and mEGFRdel19.2 readily formed lung tumors in C57BL/6 mice upon direct inoculation into the lung. By contrast, the frequency of tumor formation with mEGFR-L860R.1 cells was low with evidence of spontaneous regression in some instances. For this reason, *in vivo* evaluation of osimertinib responses will focus on mEGFRdel19.1 and mEGFRdel19.2 cells. When mice bearing orthotopic mEGFRdel19.1 and mEGFRdel19.2 tumors were submitted to daily treatment with osimertinib (5 mg/kg by oral gavage) starting 10 days after implantation, both models underwent prompt and marked tumor shrinkage (**Figure 4A and B**). Moreover, the TKI responses were durable with no evidence of tumor progression over the ∼50-day course of osimertinib treatment.

**Figure 3.**
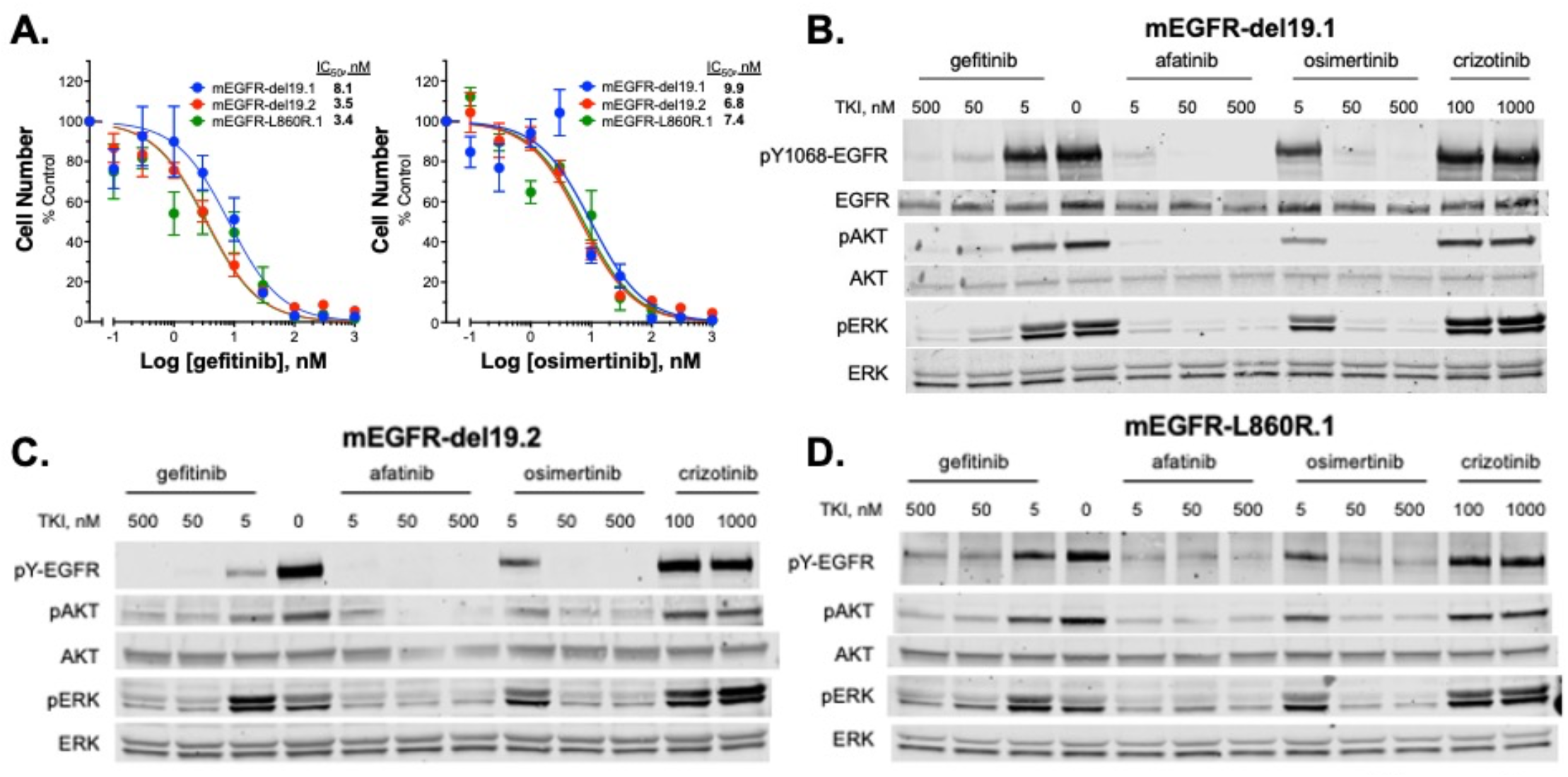
Murine EGFR-mutant LUAD cell lines are responsive to EGFR inhibitors *in vitro*. **(A)** The three EGFR-mutant murine cell lines were treated for 10 days with increasing concentrations of EGFR-specific TKIs, gefitinib or osimertinib and cell number was quantified with CyQUANT reagent. IC_50_ values were calculated with Prism 9 and presented in the graph. The data shown are the means of two independent experiments, each performed with technical triplicates. **(B-D)** The murine EGFR mutant cell lines were treated (2 hrs) with the indicated EGFR targeting TKIs (gefitinib, afatinib, osimertinib) as well as the ALK/MET-targeting TKI, crizotinib as a negative control. Cell lysates were submitted to SDS-PAGE and immunoblotted for phospho-Y1068 and total EGFR, phospho-S483 and total AKT, and phospho-Thr202/Tyr204 and total ERK.

**Figure 4.**
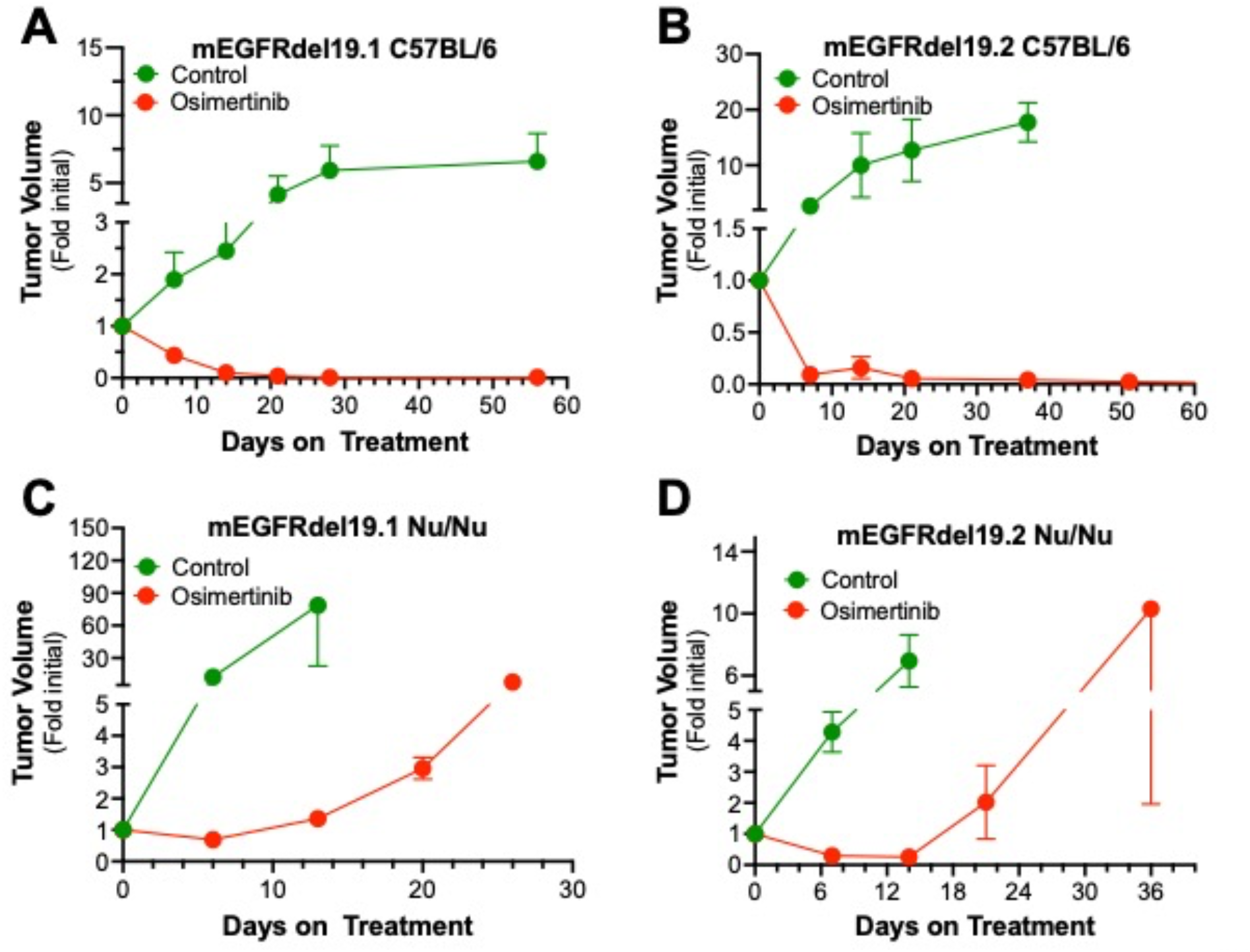
Therapeutic response of orthotopic murine EGFR tumors to osimertinib in immunocompetent and immunodeficient mice. Murine EGFRdel19.1 and del19.2 cells (500,000 cells/mouse) were injected into the left lungs of syngeneic C57BL/6 **(A, B)** or *nu/nu* **(C, D)** mice. After 10 days, the mice were imaged by *μ*CT to generate pre-treatment tumor volumes and randomized into osimertinib (5 mg/kg) or control treatment groups. The tumor-bearing mice were submitted to weekly μCT scans and the data are presented as fold of the initial pre-treatment volumes (means and SEM). For the C57BL/6 experiments, the initial tumor volumes (mean ± SEM) for the diluent and osimertinib-treated groups were 19.9 ± 4.4 mm^3^ (n=14) and 13.5 ± 2.8 mm^3^ (n=14) and 11.9 ± 4.5 mm^3^ (n=8) and 11.6 ± 3.0 mm^3^ (n=9) for mEGFRdel19.1 and mEGFRdel19.2, respectively. For the *Nu/Nu* experiments, the initial tumor volumes for the diluent and osimertinib-treated groups were 2.9 ± 0.7 mm^3^ (n=6) and 9.0 ± 1.6 mm^3^ (n=9) and 21.7 ± 6.5 mm^3^ (n=6) and 13.6 ± 4.0 mm^3^ (n=9) for mEGFRdel19.1 and mEGFRdel19.2, respectively.

### Evidence for an adaptive immunity requirement for durable osimertinib responses

Our recent studies with human EGFR mutant lung tumor biopsies (see [18] and **Fig. 1A and B**) suggest a role for host immunity in the therapeutic response to EGFR-targeting TKIs. To directly test the role for adaptive immunity, orthotopic tumors were established with mEGFRdel19.1 and mEGFRdel19.2 cells in *nu/nu* mice which lack functional T cells. As shown in **Figure 4C and D**, osimertinib treatment induced tumor shrinkage to ∼70 and 25% of initial volume in mEGFRdel19.1 and mEGFRdel19.2, respectively. In contrast to the stable therapeutic response observed in tumors propagated in C57BL/6 mice, tumor progression was observed within ∼2-3 weeks despite continuous treatment with osimertinib (**Fig. 4C and D**). Of note, propagation of human H1650 cells bearing an EGFR exon 19 deletion mutation as a flank xenograft in *nu/nu* mice revealed similar kinetics of initial tumor shrinkage and progression upon treatment with continuous osimertinib (**Suppl. Fig. S3**). These data indicate that adaptive immune cells significantly contribute to the durability of the *in vivo* TKI response in murine models driven by oncogenic EGFR.

Based on the findings in **Figure 4C** and **D** indicating a requirement for adaptive immunity for a durable osimertinib response, the intra-tumoral content of T cells was assessed by anti-CD3 immunofluorescence staining of mEGFRdel19.1 and mEGFRdel19.2 orthotopic tumors treated for 4 days with diluent or osimertinib. Representative fields from control and osimertinib-treated mEGFRdel19.2 tumors are shown in **Figure 5A** and **B**. Quantification of 3 fields from 3-4 independent tumors revealed statistically significant increases in T cell content in both mEGFRdel19.1 and mEGFRdel19.2 models treated with osimertinib. Thus, both murine EGFRdel19 tumor models exhibit baseline T cell infiltration that is further increased after 4 days of osimertinib treatment.

**Figure 5.**
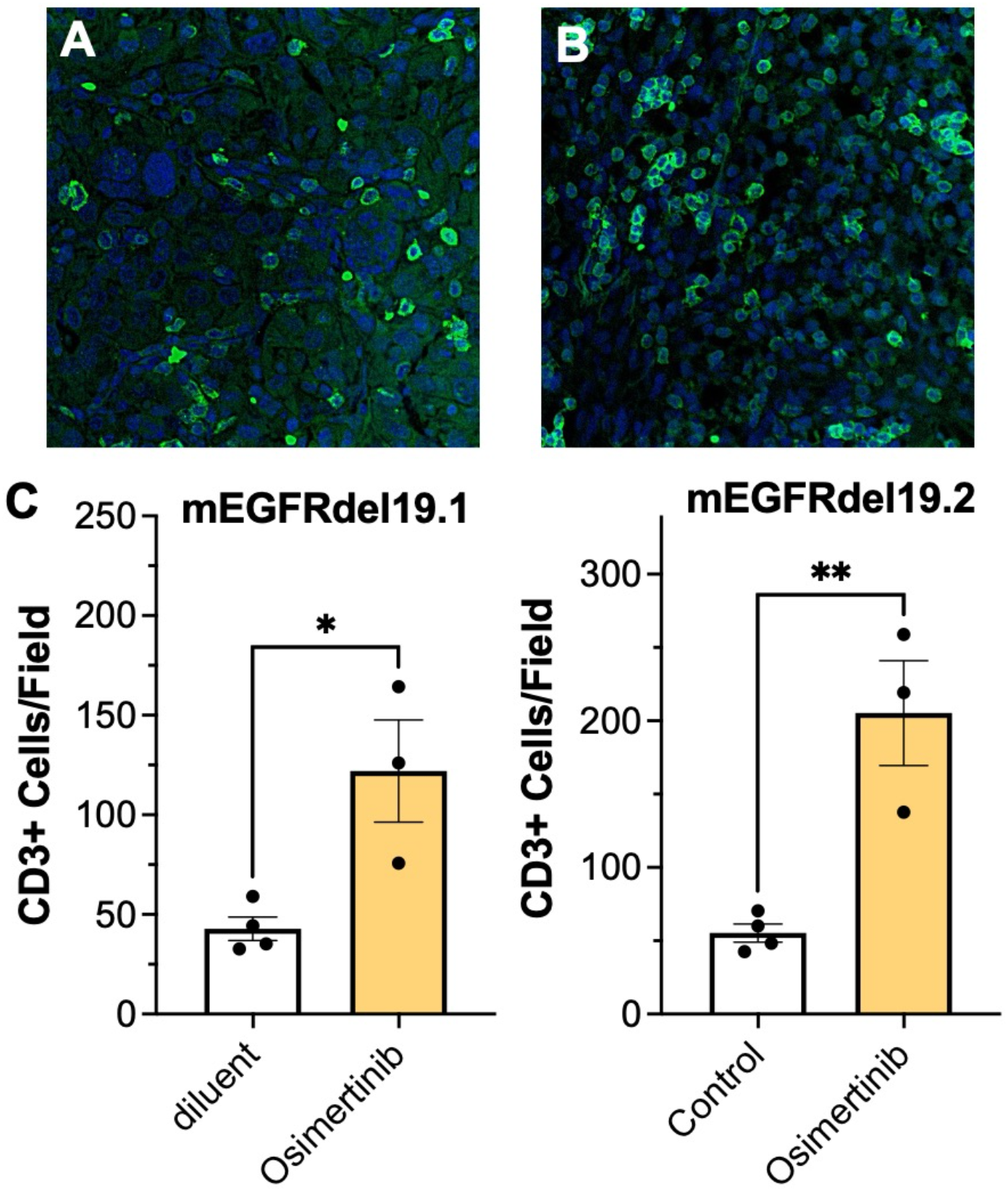
Osimertinib-treatment increases T cell content in orthotopic murine EGFRdel19 tumors. Murine EGFRdel19.1 and 19.2 cells were injected into C57BL/6 mice, tumors were permitted to establish for 14-21 days and then the mice were treated for 4 days with diluent or osimertinib. Lungs were harvested, inflated with formalin, paraffin-embedded and 5 *μ*m sections were submitted to immunofluorescence staining with anti-CD3 antibody (see Materials and Methods). CD3+ cells were quantified in 3 distinct fields for each tumor by two independent observers and the resulting values were averaged. Representative images from a mEGFRdel19.2 tumor treated with diluent (**A**) or osimertinib (**B**) are shown. The data in **C** are the means and SEM of 4 and 3 independent tumors for control and osimertinib-treated tumors, respectively, and were analyzed by student’s two-way t-tests.

## Discussion

Herein, we report the development and validation of novel EGFR mutant mouse models that recapitulate key features of human EGFR mutant lung adenocarcinoma. Furthermore, we isolated and characterized three novel murine EGFR mutant lung cancer cell lines. Combined, these GEMMs and cell lines will be valuable for dissecting the contribution of the lung TME and host immunity to therapeutic responses to TKIs that cannot be assessed with human EGFR mutant cell lines propagated in immunodeficient mice. Coupled with our orthotopic model, we were able to assess tumor shrinkage and progression in mice with intact immune systems and a lung-specific microenvironment. We were able to track individually-implanted tumors via μCT imaging to assess their dynamic responses to the 3^rd^ generation EGFR inhibitor, osimertinib. In future studies, it will be possible to molecularly engineer alter these cell lines through gene transfer-mediated overexpression or CRISPR/Cas9-mediated knockout to test specific cancer cell-derived signal pathways for roles in cancer cell-TME cross-talk and therapeutic response. In this regard, RNAseq analysis of the three murine cell lines treated *in vitro* with osimertinib (**Figure 6**) reveals marked, yet varied, regulation of chemokine and cytokine gene expression as well as putative proximally-acting transcription factor pathways (STAT1, IRF7, NFKBIA). These findings are consistent with our published findings on transcriptional reprogramming of human EGFR mutant LUAD and head and neck cancer cell lines upon treatment with EGFR-specific TKIs [18, 47]. Molecular manipulation of distinct chemokine and cytokine expression pathways that have been shown to recruit specific cell types is predicted to alter TKI responsiveness through changes in the baseline and therapy-induced immune microenvironments.

**Figure 6.**
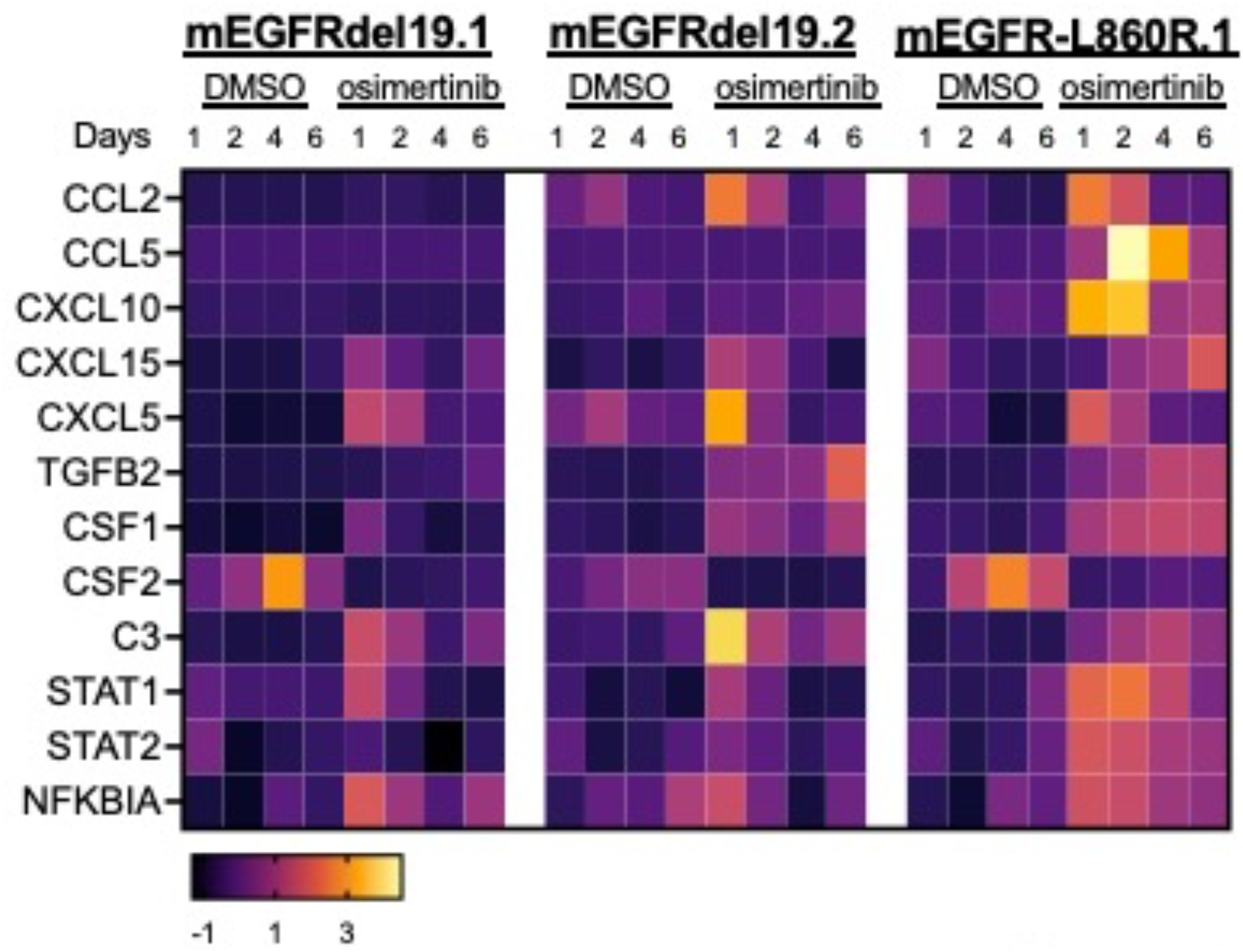
Transcriptional regulation of chemokines and cytokines in response to osimertinib treatment. The murine EGFR mutant cell lines were treated *in vitro* for 1 to 6 days with DMSO or osimertinib (100 nM) and RNA was purified and sequenced. The RNA expression values for selected genes were converted to Z-scores and presented in a heatmap format.

Using these murine EGFR mutant cell lines, we present strong evidence for adaptive immunity as a requirement for durable inhibition of orthotopic tumors by osimertinib (**Figure 4**). In this regard, our present study adds to a growing body of evidence supporting host immunity as a contributor to therapeutic efficacy of oncogene-targeted agents [20-27, 48]. We previously reported that the degree of induction of the IFNγ Hallmark response as well as a T cell signature [49] in on-treatment biopsies associate with the duration of therapeutic benefit in patients bearing EGFR mutant lung cancers [18]. Herein, we demonstrate that the magnitude of two additional gene expression signatures previously shown to predict responsiveness to anti-PD-1-based immune therapy [42, 43] also associate with longer time to progression on EGFR TKIs (**Figure 1**). Thus, there is ample evidence for immune engagement by human EGFR mutant lung cancers despite their lack of sensitivity to anti-PD-1 therapies [44-46], a feature that is shared by orthotopic tumors derived from the murine EGFR mutant cell lines (data not shown). While the aforementioned evidence for immune cell engagement by EGFR mutant lung cancers provides rationale for combining EGFR-specific TKIs and anti-PD-1 immunotherapy, clinical trials testing these combinations were terminated early due to toxicities [50, 51]. In sum, the published and present findings suggest that host immunity is critical for durability of TKI response, but perhaps is regulated in a manner distinct from that associated with response to anti-PD-1 immunotherapy. A recent study by Tang et al [25] exploring sensitivity to SHP2 inhibitors revealed a dominant role for immunosuppressive myeloid cell types. Blockade of myeloid cell recruitment by CXCR1/2 inhibitors markedly enhanced T cell function and overall therapeutic response. Moreover, a study by Maynard et al demonstrated accumulation of immune suppressive cell types after longer times of TKI treatment in EGFR and ALK-driven lung tumors [[19]. Thus, alternative means to boost cytotoxic T cell activity besides anti-PD-1 agents may be required to enhance the contribution of adaptive immunity to TKI responsiveness in EGFR mutant lung cancers. These transplantable murine EGFR cell lines should prove invaluable for a deeper interrogation of the immune microenvironment in this oncogene-defined LUAD subset with the goal of identifying a novel combination therapy that bolsters adaptive immunity in conjunction with EGFR-specific TKIs to prolong the duration of therapeutic benefit presently experienced by patients.

A simple interpretation of the rapid progression of murine EGFR mutant tumors propagated in the absence of functional adaptive immunity (**Figure 4C, D**) is that immune surveillance by specific T cell and/or B cell populations serves as a break on tumor progression driven by acquired resistance in the setting of TKI treatment. Thus, variation in the efficacy of immune surveillance amongst EGFR mutant lung cancer patients may account for the observed range in time to progression. While not explored in this study, it is interesting to consider the mutational and/or differentiation status of the murine EGFR mutant tumors that rapidly emerge in *nu/nu* mice. The spectrum of EGFR kinase domain mutations that confer osimertinib resistance has been defined in human EGFR mutant LUAD [13, 14], but the rapidity with which the emergence of resistance occurs seems more consistent with transcription-dependent phenotype switching [52] or engagement of bypass signaling pathways. It is possible that initial shrinkage of the tumors with TKI treatment alters the balance between T cells and cancer cells as proposed in the immunoediting model of cancer [53] and prevents the ability of residual cancer cells undergoing bypass signaling to proliferate. Preliminary *in vitro* RNAseq analysis of the murine EGFR mutant cell lines reveals osimertinib-induction of HGF-MET and FGFR pathways as candidates (data not shown), but it is uncertain how the lack of adaptive immunity permits these bypass signaling pathways to be functionally unleashed. Future studies will be directed at characterizing the progressing tumors in the immunodeficient hosts.

## Author Contributions

Concept and experimental design: EKK, ATL, ETC, CJH, MWE, RAN, LEH

Developed methodology: EKK, ATL, TKH, CJH, MWE, RAN, LEH

Performed experiments and acquired data: EKK, ATL, TKH, DTM, TTN, AN, LEH

Analyzed and interpreted data: EKK, ATL, TKH, TTN, AN, DTM, RAN, LEH

Prepared figures and drafted manuscript: EKK, RAN, LEH

Edited and revised manuscripts: EKK, RAN, LEH

## Acknowledgements

We would like to acknowledge Dr. Jennifer Matsuda and the Mouse Genetics Core at National Jewish Health for generating the murine EGFR^del19^ and EGFR^L860R^ GEMMs. The RNA sequencing was performed by the Genomics shared resource within the University of Colorado Cancer Center.

