## Supplementary Information for "Novel EGFR-Mutant Mouse Models of Lung Adenocarcinoma Reveal Adaptive Immunity Requirement for Durable Osimertinib Response"

**Supplementary Table S1**

**Supplementary Figures S1-S3**

| Supplementary Table S1. Quantification of Lesions in Murine EGFR Transgenic Mice | | | | | | |
|---|---|---|---|---|---|---|
| Genotype | Weeks post AdCre | Sex | hyperplasias | AAH | adenomas | adenocarcinomas |
| $Egfr^{+/del19}$; $p53^{+/+}$ | 14 weeks | F | 0 | 3 | 1 | 6 |
| $Egfr^{+/del19}$;$p53^{+/f}$ | ~5.5 weeks | F | 6 | 0 | 0 | 0 |
| $Egfr^{+/L860R}$;$p53^{+/f}$ | 11 weeks | M | 0 | 9 | 3 | 0 |
| $Egfr^{+/del19}$;$p53^{+/+}$ | ~13 weeks | M | 3 | 20 | 12 | 2 |
| $Egfr^{+/L860R}$;$p53^{+/f}$ | ~11 weeks | M | 4 | 7 | 2 | 0 |
| $Egfr^{L860R/L860R}$;$p53^{+/+}$ | 14 weeks | F | 0 | 1 | 1 | 4 |
| $Egfr^{+/L860R}$;$p53^{+/f}$ | 8 weeks | F | 0 | 0 | 0 | 0 |
| $Egfr^{+/del19}$;$p53^{+/f}$ | 8 weeks | F | 2 | 17 | 16 | 4 |
| $Egfr^{+/L860R}$;$p53^{+/f}$ | 11 weeks | M | 0 | 0 | 0 | 0 |
| $Egfr^{+/del19;}p53^{+/f}$ | 11 weeks | F | 1 | 51 | 29 | 0 |

Mice bearing the indicated transgene were administered adeno-Cre virus by intratracheal instillation and after the indicated number of weeks, euthanized. The lungs were harvested, inflated and formalin-fixed. Following paraffin embedding, sections were stained with hematoxylin and and the numbers of hyperplasias, atypical adenomatous hyperplasias (AAH), adenomas and adenocarcinomas was quantified by a pathologist.

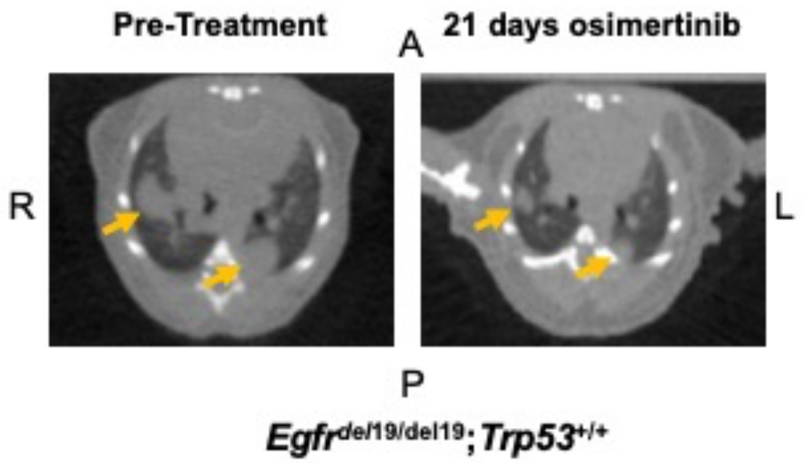


### Supplementary Figure S1

**Supplemental Figure S1. Primary tumors emerging in an EGFR-mutant GEMM are responsive to osimertinib.** Approximately 15 weeks after intratracheal AdCre instillation, the indicated mEGFR-del19$^{+/+}$/Trp53$^{+/+}$ mouse was imaged by μCT which revealed multifocal tumor formation (see arrows). Daily oral gavage with 5 mg/kg of osimertinib was initiated and continued for 3 weeks. Repeat μCT imaging revealed tumor shrinkage of the lesions. A=anterior, P=posterior, R=right, L=left.

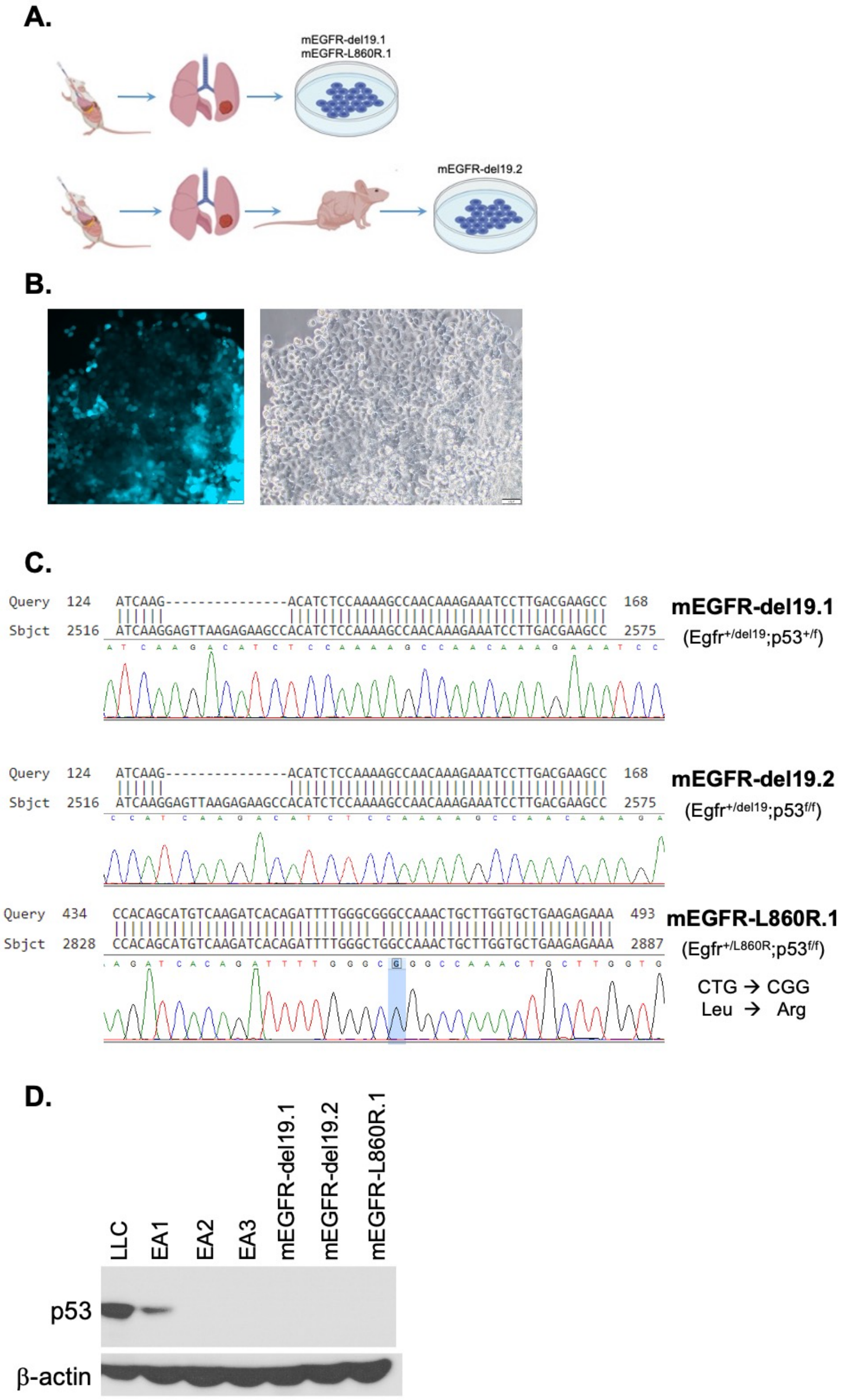


**Supplementary Figure S2**

**Supplemental Figure S2. Establishment and characterization of cell lines from Egfr$^{del19}$ and Egfr$^{L860R}$ lung tumors. (A)** Schematic of strategies used to generate cell lines. Adeno-cre virus was instilled into Egfr$^{del19}$ and Egfr$^{L860R}$ mice and tumors were allowed to establish. Individual tumors were harvested from mice, dissociated, and grown on plastic in order to develop 2 cell lines (mEGFR-del19.1, mEGFR-L860R.1). To create the mEGFR-del19.2 cell line, a tumor was dissected, cut into approximately 1 mm$^3$ pieces and implanted into the flanks of *nu/nu* mice. Once tumors established, they were dissociated and cultured on plastic to create the cell line. **(B)** Cerulean blue fluorescence (left) and phase contrast (right) images of the mEGFR-del19.2 cell line. Scale bars = 50 μm, magnification is 100x. **(C)** Sanger sequencing of the Egfr PCR amplicon indicating the predicted Egfr mutations. **(D)** Cell extracts were prepared from mEGFRdel19.1, 19.2 and L860R.1 as well as 3 distinct murine Eml4-Alk positive cell lines (EA1, TP53 wild-type, EA2 and EA3, TP53 null) and KRAS-G12C mutant LLC cells bearing mutated TP53. The extracts were submitted to SDS-PAGE and immunoblotted for murine TP53. The filter was stripped and re-probed for β-actin as a loading control.

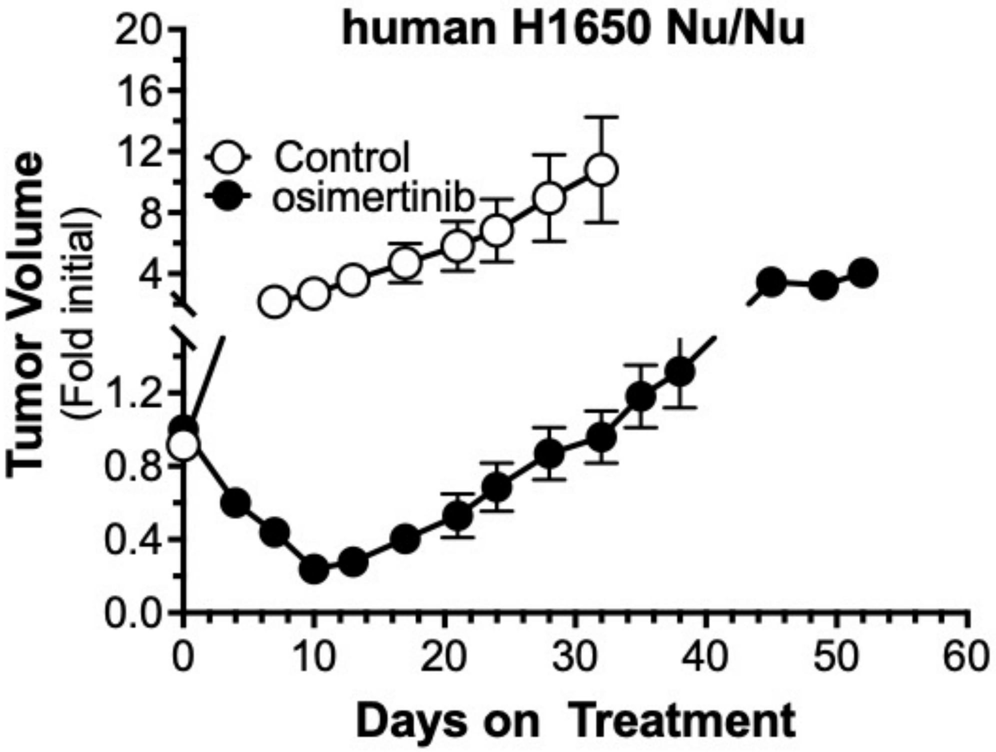


**Supplementary Figure S3**

**Supplemental Figure S3. Kinetics of tumor shrinkage and progression in human H1650 xenografts propagated in nu/nu mice.** Human H1650 cells bearing an EGFR exon 19 deletion were inoculated into the flanks of *nu/nu* mice. When the tumors reached ~200 $mm^3$, the mice were randomized to diluent control or 5 mg/kg osimertinib delivered by oral gavage daily. Tumor volumes were measured every three days with calipers and presented as fold of initial pre-treatment volumes. The data are the means and SEM with 12 mice per group.
